# Transcriptomic FOLFIRINOX sensitivity signatures stratify overall survival and reveal directional reclassification after neoadjuvant FOLFIRINOX in borderline resectable and locally advanced pancreatic cancer

**DOI:** 10.64898/2026.09.07.749857

**Authors:** Clémentine Gaucher, Muge Kaya, Jonathan Garnier, Anna Barthélémy, Philippe Rochigneux, Flora Poizat, Marion Rubis, Pauline Moussard, Julie Roques, Nicolas Fraunhoffer, Nelson Dusetti, Brice Chanez

**Affiliations:** Aix-Marseille University, Inserm, CNRS, Institut Paoli-Calmettes, Cancer Research Centre of Marseille (CRCM), Marseille, France; Department of Surgical Oncology, Institut Paoli-Calmettes, Marseille, France; Department of Pathology, Institut Paoli-Calmettes, Marseille, France; Experimental Histopathology ICEP Platform, Centre de Recherche en Cancérologie de Marseille (CRCM), Aix-Marseille Université, CNRS, INSERM, Institut Paoli-Calmettes, Marseille, France; Department of Medical Oncology, Institut Paoli-Calmettes, Marseille, France

**Keywords:** Pancreatic ductal adenocarcinoma, FOLFIRINOX, transcriptomic signatures, neoadjuvant chemotherapy, treatment response, precision oncology

## Abstract

**Background:** Transcriptomic FOLFIRINOX-component sensitivity signatures have been clinically evaluated in resected and advanced PDAC, but their relevance and longitudinal stability in the neoadjuvant BR/LA setting are unknown.

**Patients and methods:** We retrospectively studied 77 patients with borderline resectable or locally advanced PDAC treated with neoadjuvant FOLFIRINOX followed by resection. Pretreatment sensitivity to 5-fluorouracil, oxaliplatin and irinotecan was determined using previously developed locked transcriptomic classifiers and integrated into a regimen-level FOLFIRINOX classification. The primary molecular cohort comprised 53 patients whose pretreatment biopsies had ≥10% tumour cellularity. Cox models assessed associations with survival. Longitudinal changes were assessed in 35 paired tumours.

**Results:** Thirty-one pretreatment tumours were classified as FOLFIRINOX-sensitive (FFX-Sens), and 22 were not classified as FOLFIRINOX-sensitive (FFX-Res). FFX-Sens status was associated with longer overall survival (OS; hazard ratio [HR] 0.48, 95% confidence interval [CI] 0.25-0.93; p=0.029). In the complete-case model adjusted for baseline carbohydrate antigen 19-9 (CA19-9) and tumour size (n=50), the association with OS persisted (adjusted HR 0.47, 95% CI 0.23-0.96; p=0.037), whereas the adjusted association with disease-free survival was not statistically significant (adjusted HR 0.55, 95% CI 0.28-1.10; p=0.090). Among paired tumours, 15 changed from FFX-Sens to FFX-Res and four in the opposite direction (exact McNemar p=0.019). The OS association also persisted after adjustment for postoperative pathological factors (adjusted HR 0.40; p=0.015).

**Conclusions:** Pretreatment FOLFIRINOX sensitivity was associated with OS under neoadjuvant FOLFIRINOX, while matched pretreatment and residual-tumour analyses revealed significant directional reclassification after treatment. These findings extend the clinical evaluation of validated drug-specific FOLFIRINOX sensitivity classifiers to the neoadjuvant setting and provide a longitudinal assessment of their stability during treatment. They support prospective evaluation of both pretreatment stratification and molecular reassessment of residual disease.

**Highlights:**

- Pretreatment FOLFIRINOX sensitivity was associated with overall survival.
- The association persisted after adjustment for CA19-9 and tumour size.
- Residual tumours show directional reclassification after FOLFIRINOX.
- The overall-survival association persisted after pathological adjustment.

## Introduction

Pancreatic ductal adenocarcinoma (PDAC) remains one of the most lethal solid malignancies, with systemic relapse occurring frequently even in patients presenting with apparently localised disease. In borderline resectable and locally advanced PDAC (BR/LA), neoadjuvant chemotherapy has become a standard of care, enabling early treatment of occult metastatic disease and providing an opportunity to assess tumour behaviour before surgery [1]. The best regimen and duration of induction chemotherapy nevertheless remain largely empirical [2–5]. FOLFIRINOX and gemcitabine-based regimens are both used, but substantial interpatient heterogeneity exists in treatment efficacy and toxicity, and no validated tumour-specific biomarker currently guides regimen selection. This issue is particularly relevant in the potentially curative setting, where progression during induction chemotherapy may compromise surgery and survival [1].

Transcriptomic profiling has established PDAC as a biologically heterogeneous disease. Molecular classifications initially identified distinct tumour subtypes [6–8], broadly converging on classical and basal-like phenotypes, with the latter generally associated with more aggressive disease and reduced benefit from conventional chemotherapy [9–11]. These classifications provide important biological and prognostic information but were not specifically designed to estimate tumour sensitivity to individual cytotoxic drugs. To address this limitation, we developed drug-specific transcriptomic signatures derived from functionally characterised patient-derived PDAC models. GemPred was initially developed to predict gemcitabine sensitivity in resected PDAC and subsequently validated in the randomised PRODIGE-24/CCTG PA.6 cohort [12,13]. *GemCore* extended this strategy to advanced disease and limited biopsy material [14], and the same model-based approach was subsequently applied to predict sensitivity to 5-fluorouracil, oxaliplatin and irinotecan, generating the *5FUCore*, *OxaCore* and *IriCore* signatures [15]. In patients with advanced PDAC, survival progressively improved with the number of FOLFIRINOX components to which tumours were predicted to be sensitive [15]. These drug-specific predictors were subsequently incorporated into broader transcriptomic approaches evaluated in independent clinical settings [16–18]. However, previous clinical evaluations of these drug-sensitivity classifiers were performed in resected/adjuvant or advanced/metastatic PDAC; their clinical relevance when applied to pretreatment biopsies before neoadjuvant therapy in BR/LA PDAC has not been reported. This represents an important clinical gap. In BR/LA disease, initial systemic therapy is given not only to control occult systemic disease but also to preserve or achieve resectability. Failure to control the tumour during this interval may eliminate the opportunity for curative-intent resection, while regimen selection remains largely empirical.

The neoadjuvant setting provides a clinically relevant opportunity to evaluate this strategy. Pretreatment pancreatic biopsies are obtained before exposure to systemic therapy and could therefore provide molecular information to inform treatment decisions. Surgical resection after neoadjuvant treatment provides access to the residual tumour population that has persisted despite several cycles of chemotherapy. Paired analysis of these specimens can therefore address two complementary questions: whether a pretreatment drug-sensitivity profile is associated with subsequent clinical outcome, and whether that molecular sensitivity state persists in residual tumour following treatment. Although longitudinal transcriptomic studies have described molecular changes after neoadjuvant therapy, the stability of validated drug-specific FOLFIRINOX sensitivity classifiers has not, to our knowledge, been examined in matched pretreatment and residual-tumour samples. Reassessment of residual disease could therefore characterise the molecular state of the tumour that persists after treatment and explore whether post-treatment classification might inform subsequent therapeutic decisions; this potential use remains untested.

Here, we evaluated previously developed transcriptomic FOLFIRINOX sensitivity signatures in patients with PDAC treated with neoadjuvant FOLFIRINOX followed by surgical resection. We assessed whether the pretreatment FOLFIRINOX sensitivity profile was associated with survival and whether its association with outcome persisted after accounting for major baseline clinical factors. We also examined whether pretreatment molecular information complemented postoperative pathological assessment. Using paired pretreatment biopsies and residual surgical tumours, we investigated longitudinal changes in FOLFIRINOX sensitivity and its individual drug components. Because all patients received FOLFIRINOX before surgery and no alternative neoadjuvant treatment arm was available, the study evaluates outcome stratification under FOLFIRINOX rather than formally establishing treatment-specific predictive value. To our knowledge, this is the first study to evaluate these validated FOLFIRINOX-component drug-sensitivity classifiers in the neoadjuvant BR/LA setting and the first to examine their longitudinal stability in matched pretreatment and residual-tumour specimens.

## Methods

### Study design and patient population

This was a retrospective, single-centre, observational cohort study of analytical (explanatory) design, evaluating the association between pretreatment transcriptomic FOLFIRINOX sensitivity and survival outcomes after neoadjuvant FOLFIRINOX. This translational study required close collaboration between the surgical oncology, pathology, medical oncology and bioinformatics teams at Institut Paoli-Calmettes and the Cancer Research Centre of Marseille. Patients were identified from a retrospective surgical cohort of 141 patients with BR/LA PDAC who underwent pancreatic resection at Institut Paoli-Calmettes between June 2014 and October 2022. After exclusion of 28 patients who did not meet the prespecified clinical eligibility criteria, 113 patients remained eligible. The exclusion criteria were performance status ≥2 at diagnosis; no induction treatment; induction treatment other than FOLFIRINOX; chemoradiotherapy; histology other than adenocarcinoma; metastatic disease in the surgical specimen; recurrence or death within 3 months after surgery; and refusal of data use. Usable pretreatment biopsy material was available for 60 patients; 17 additional patients with usable resection specimens but no pretreatment biopsy were included for analyses of surgical material, yielding an overall translational cohort of 77 patients (Figure 1).

**Figure 1.**
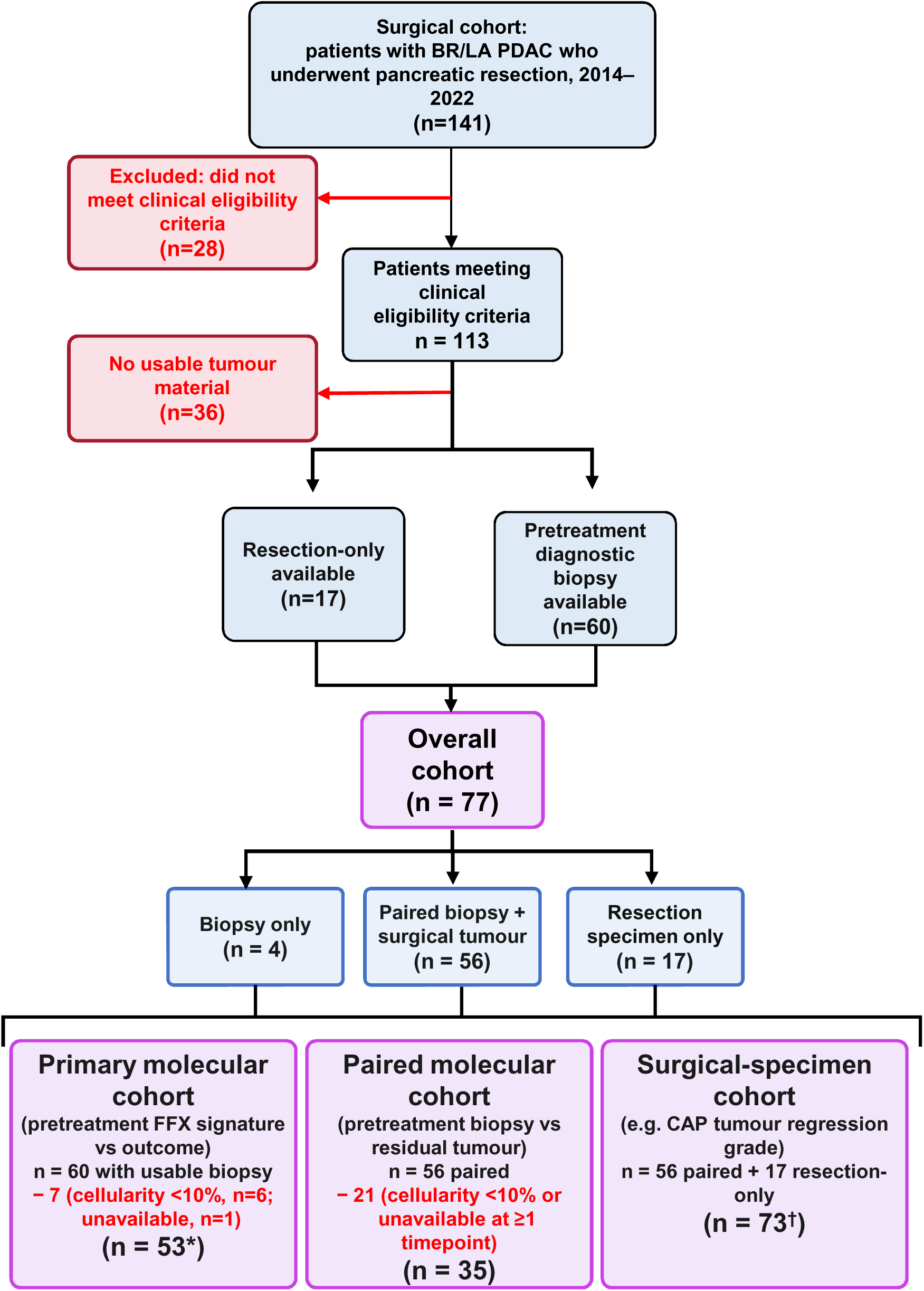
Study flow and definition of molecular analysis cohorts. The source surgical cohort comprised 141 patients with BR/LA PDAC who underwent pancreatic resection between June 2014 and October 2022. After exclusion of 28 patients who did not meet the clinical eligibility criteria, 113 patients remained. Pretreatment diagnostic biopsy material was available for 60 patients. Among the 53 patients without a usable pretreatment biopsy, 17 with resection-only FFPE material were included and 36 had no usable material for translational analyses, yielding an overall translational cohort of 77 patients. Seven patients were excluded from the primary molecular analysis because pretreatment biopsy tumour cellularity was <10% (n=6) or unavailable (n=1), resulting in a primary molecular cohort of 53 patients. Surgical tissue blocks could not be retrieved for four patients, leaving 56 patients with paired pretreatment biopsy and surgical specimens. After exclusion of 21 patients with tumour cellularity <10% or unavailable in at least one specimen, 35 patients comprised the paired molecular cohort. The paired molecular cohort was a subset of the primary molecular cohort. Because assessment of the operated tumour (e.g. pathological response) does not require a pretreatment biopsy, both patients with paired biopsy and surgical specimens and patients with resection-only material provided usable tumour tissue, yielding 73 patients (56 paired plus 17 resection-only) for analyses restricted to the surgical specimen. * Complete-case multivariable analyses were restricted to 50 patients. For baseline models, CA19-9 and/or tumour size data were missing in 3 patients from the primary molecular cohort; for postoperative models, CAP tumour regression grade was unavailable in 3 patients. † Operated-tumour analyses used the surgical-specimen cohort (n=73), but CAP tumour regression grade was evaluable in 72 patients. BR/LA, borderline resectable/locally advanced; FFPE, formalin-fixed, paraffin-embedded; PDAC, pancreatic ductal adenocarcinoma FFX, FOLFIRINOX; CAP, College of American Pathologists.

Disease extent was reviewed by the multidisciplinary tumour board and classified according to the anatomical, biological and conditional criteria of the 2017 international consensus [19]. All patients received at least two cycles of neoadjuvant FOLFIRINOX before surgery. All patients received at least two cycles of neoadjuvant FOLFIRINOX before surgery. Treatment was based on the original FOLFIRINOX regimen [20], consisted of oxaliplatin 85 mg/m², irinotecan 180 mg/m², folinic acid 200 mg/m2 and fluorouracil 2400 mg/m² administered as a 46-h continuous infusion without a fluorouracil bolus.

The study population was defined by successful completion of the neoadjuvant-to-surgery pathway. Patients who progressed, died or became unresectable during neoadjuvant treatment were therefore not represented in this surgical cohort. Clinical, treatment, pathological and survival data were retrospectively collected from institutional records. Postoperative chemotherapy was administered according to clinical practice and was not randomised. Clinical data were compiled in a dedicated database from patients’ electronic health records.

### Ethics approval and consent

Patients were identified from the prospective single-centre CHIRPAN pancreatic surgery database (NCT02871336), sponsored by Institut Paoli-Calmettes. The CHIRPAN protocol received a favourable opinion from the Research Ethics Committee on 31 May 2021 (N°21037-59833), and the database had been authorised by the French data-protection authority, the Commission nationale de l’informatique et des libertés (CNIL), on 14 October 2014. The present translational study was approved by the institutional review board in 2022 (NEOPRONOX, IPC 2022-052). Patients received written and oral information, and their non-opposition to participation and to the use of their clinical and molecular data for research was documented in the medical record, in accordance with the Declaration of Helsinki.

### Tumour samples, RNA extraction and sequencing

Pretreatment tumour material was obtained from formalin-fixed, paraffin-embedded (FFPE) diagnostic pancreatic biopsy specimens collected by endoscopic ultrasound-guided sampling. Post-treatment tumour material was obtained from surgical resection specimens after FOLFIRINOX. FFPE material was processed according to the previously published workflow used for the development and clinical evaluation of the PDAC chemotherapy-sensitivity signatures [15].

Briefly, 10-µm sections were cut from each FFPE block and macrodissected to enrich for neoplastic tissue. Total RNA was extracted using the RNeasy FFPE kit (Qiagen) according to the manufacturer’s instructions. RNA libraries were prepared using the QuantSeq 3′ mRNA-Seq kit (Lexogen, Vienna, Austria), as previously described [15]. Sequencing was performed using an Illumina NovaSeq 6000 system, targeting approximately 25 million reads per sample. Reads were aligned to the human hg38 reference genome with the Rsubread R package (version 2.24.0). Raw counts were normalised using the trimmed mean of M-values method (TMM; edgeR version 3.44.0). Neoplastic cellularity was estimated by a pathologist as the percentage of malignant epithelial cells within the analysed material.

### Transcriptomic FOLFIRINOX sensitivity classification

Sensitivity to 5-fluorouracil, oxaliplatin and irinotecan was determined using the previously developed and locked *5FUCore*, *OxaCore* and *IriCore* transcriptomic classifiers [15]. The classifiers were applied without retraining, parameter optimisation or cohort-specific recalibration.

A regimen-level FOLFIRINOX classification was derived from the three individual drug-sensitivity calls. Tumours predicted to be sensitive to at least two of the three FOLFIRINOX components were classified as FOLFIRINOX-sensitive (FFX-Sens), whereas tumours predicted to be sensitive to zero or one component were classified as not FOLFIRINOX-sensitive (FFX-Res) [15]. Based on previous clinical applications of this transcriptomic workflow, in which a 10% neoplastic-cellularity threshold was used to define suitability of FFPE samples for transcriptomic analysis [14,15], pretreatment biopsies with tumour cellularity ≥10% were considered molecularly evaluable for the primary analysis.

### Pathological assessment

Pathological response was evaluated in the surgical specimen using the College of American Pathologists (CAP) tumour regression grading system [21]. Post-treatment nodal status was classified as ypN0, ypN1 or ypN2. Resection margins were classified as R0 when tumour clearance was ≥1 mm and R1 when clearance was <1 mm [22]. CAP tumour regression grades 0-1 were considered a major pathological response.

### Longitudinal molecular analyses

Longitudinal transcriptomic analyses were restricted to patients with paired pretreatment biopsy and residual surgical tumour specimens with neoplastic cellularity ≥10% at both time points. This paired molecular cohort comprised 35 patients. Changes in binary FOLFIRINOX sensitivity classification between pretreatment and residual tumour were assessed using the exact McNemar test. Changes in *5FUCore*, *OxaCore* and *IriCore* classifications were additionally examined separately to determine the contribution of each drug-specific signature to longitudinal reclassification. *GemCore* classification was also examined in the same paired samples as an exploratory negative control because gemcitabine is not a component of FOLFIRINOX [14].

### Exploratory molecular phenotype analyses

To place longitudinal changes in drug sensitivity in the broader context of PDAC molecular phenotype, the pancreatic adenocarcinoma molecular gradient (PAMG) and Purity Independent Subtyping of Tumors (PurIST) classifiers were analysed as exploratory secondary measures. PurIST classifications were generated according to the published single-sample implementation [10], and PAMG scores according to the previously described molecular-gradient framework [23]. PAMG scores were compared between paired pretreatment and residual-tumour samples using the paired Wilcoxon signed-rank test, both in all patients with paired measurements and after restriction to specimens with tumour cellularity ≥10% at both time points. Changes in PurIST classical/basal classification were evaluated in the paired molecular cohort using the exact McNemar test.

### Survival endpoints and statistical analysis

Overall survival (OS) was calculated from the date of surgery to death from any cause, with patients alive at last follow-up censored at that date. Disease-free survival (DFS) was calculated from the date of surgery to the first occurrence of oncological recurrence or death from any cause, whichever occurred first, with event-free patients censored at last follow-up. Continuous variables are reported as medians with interquartile ranges (IQRs), and categorical variables as numbers and percentages. Between-group comparisons in Table 1 used the Wilcoxon rank-sum test for continuous variables and Fisher’s exact test for categorical variables; these comparisons were exploratory. Median follow-up was estimated using the reverse Kaplan-Meier method. Survival distributions were estimated using the Kaplan-Meier method and compared using log-rank tests. Cox proportional-hazards models were used to estimate hazard ratios (HRs) and 95% confidence intervals (CIs). The proportional-hazards assumption was assessed using Schoenfeld residuals and showed no material violation. The primary survival analysis compared pretreatment FFX-Sens with FFX-Res in the primary molecular cohort. Associations between baseline clinical variables and OS or DFS were additionally assessed using univariable Cox models. Evaluated variables were FFX classification, age (>65 versus ≤65 years), sex (male versus female), performance status (1 versus 0), disease extent (locally advanced versus borderline resectable), primary tumour location (other versus head), baseline CA19-9 (≥500 versus <500 U/mL) and baseline tumour size (>40 versus ≤40 mm).

**Table 1.**
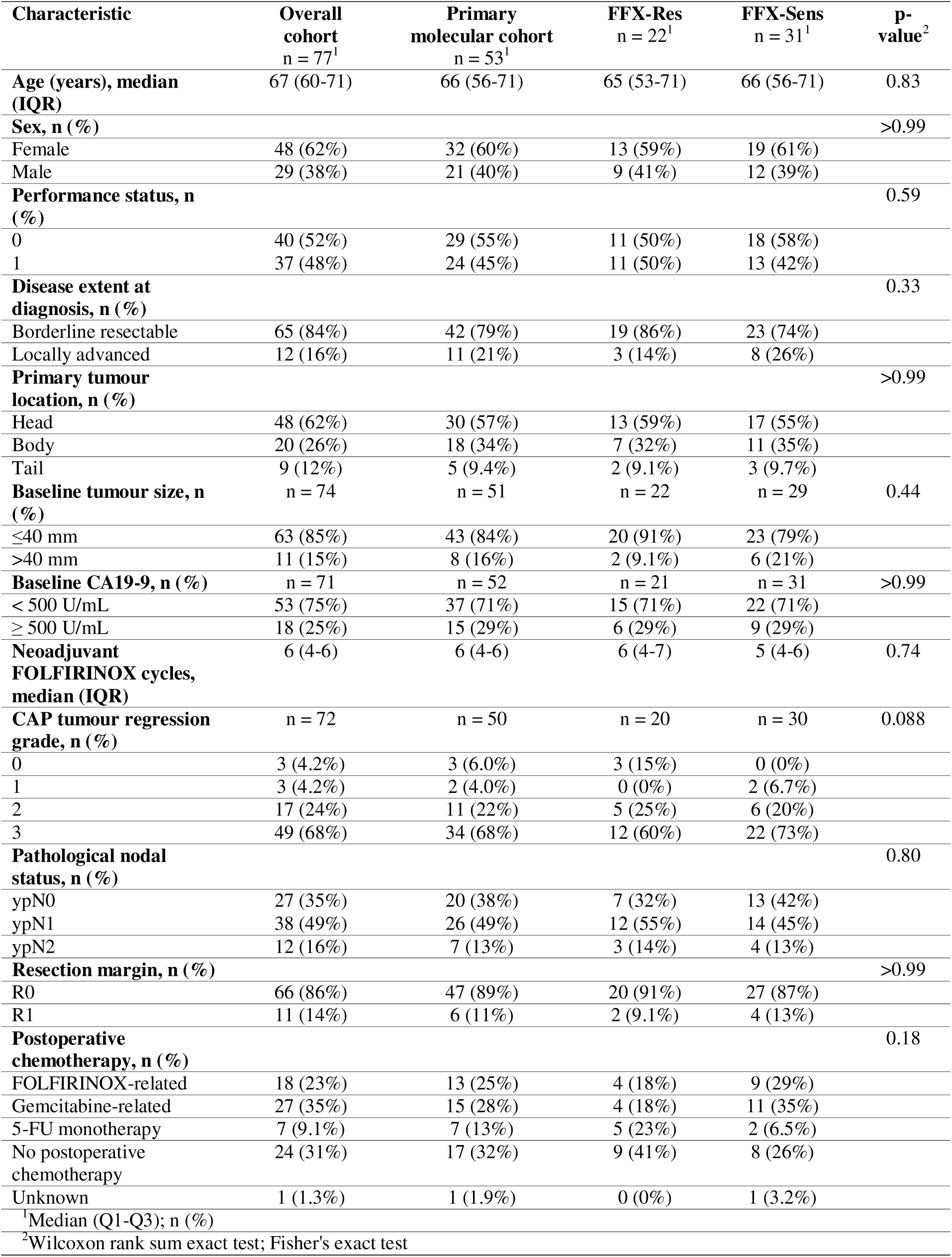
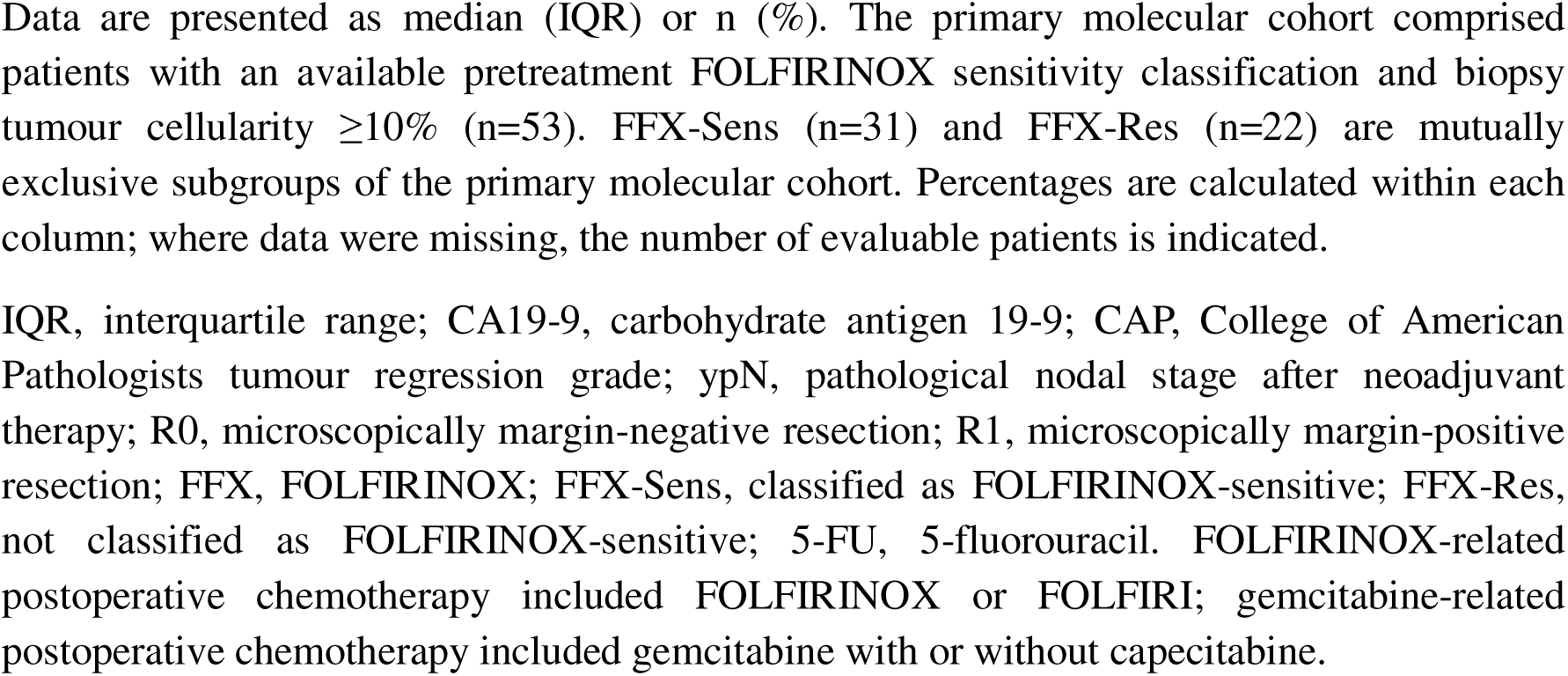
Clinicopathological characteristics of the study population.

| Characteristic | Overall cohort<br>n = 77<sup>1</sup> | Primary molecular cohort<br>n = 53<sup>1</sup> | FFX-Res<br>n = 22<sup>1</sup> | FFX-Sens<br>n = 31<sup>1</sup> | p-value<sup>2</sup> |
| --- | --- | --- | --- | --- | --- |
| **Age (years), median (IQR)** | 67 (60-71) | 66 (56-71) | 65 (53-71) | 66 (56-71) | 0.83 |
| **Sex, n (%)** |  |  |  |  | >0.99 |
| Female | 48 (62%) | 32 (60%) | 13 (59%) | 19 (61%) |  |
| Male | 29 (38%) | 21 (40%) | 9 (41%) | 12 (39%) |  |
| **Performance status, n (%)** |  |  |  |  | 0.59 |
| 0 | 40 (52%) | 29 (55%) | 11 (50%) | 18 (58%) |  |
| 1 | 37 (48%) | 24 (45%) | 11 (50%) | 13 (42%) |  |
| **Disease extent at diagnosis, n (%)** |  |  |  |  | 0.33 |
| Borderline resectable | 65 (84%) | 42 (79%) | 19 (86%) | 23 (74%) |  |
| Locally advanced | 12 (16%) | 11 (21%) | 3 (14%) | 8 (26%) |  |
| **Primary tumour location, n (%)** |  |  |  |  | >0.99 |
| Head | 48 (62%) | 30 (57%) | 13 (59%) | 17 (55%) |  |
| Body | 20 (26%) | 18 (34%) | 7 (32%) | 11 (35%) |  |
| Tail | 9 (12%) | 5 (9.4%) | 2 (9.1%) | 3 (9.7%) |  |
| **Baseline tumour size, n (%)** | n = 74 | n = 51 | n = 22 | n = 29 | 0.44 |
| ≤40 mm | 63 (85%) | 43 (84%) | 20 (91%) | 23 (79%) |  |
| >40 mm | 11 (15%) | 8 (16%) | 2 (9.1%) | 6 (21%) |  |
| **Baseline CA19-9, n (%)** | n = 71 | n = 52 | n = 21 | n = 31 | >0.99 |
| < 500 U/mL | 53 (75%) | 37 (71%) | 15 (71%) | 22 (71%) |  |
| ≥ 500 U/mL | 18 (25%) | 15 (29%) | 6 (29%) | 9 (29%) |  |
| **Neoadjuvant FOLFIRINOX cycles, median (IQR)** | 6 (4-6) | 6 (4-6) | 6 (4-7) | 5 (4-6) | 0.74 |
| **CAP tumour regression grade, n (%)** | n = 72 | n = 50 | n = 20 | n = 30 | 0.088 |
| 0 | 3 (4.2%) | 3 (6.0%) | 3 (15%) | 0 (0%) |  |
| 1 | 3 (4.2%) | 2 (4.0%) | 0 (0%) | 2 (6.7%) |  |
| 2 | 17 (24%) | 11 (22%) | 5 (25%) | 6 (20%) |  |
| 3 | 49 (68%) | 34 (68%) | 12 (60%) | 22 (73%) |  |
| **Pathological nodal status, n (%)** |  |  |  |  | 0.80 |
| ypN0 | 27 (35%) | 20 (38%) | 7 (32%) | 13 (42%) |  |
| ypN1 | 38 (49%) | 26 (49%) | 12 (55%) | 14 (45%) |  |
| ypN2 | 12 (16%) | 7 (13%) | 3 (14%) | 4 (13%) |  |
| **Resection margin, n (%)** |  |  |  |  | >0.99 |
| R0 | 66 (86%) | 47 (89%) | 20 (91%) | 27 (87%) |  |
| R1 | 11 (14%) | 6 (11%) | 2 (9.1%) | 4 (13%) |  |
| **Postoperative chemotherapy, n (%)** |  |  |  |  | 0.18 |
| FOLFIRINOX-related | 18 (23%) | 13 (25%) | 4 (18%) | 9 (29%) |  |
| Gemcitabine-related | 27 (35%) | 15 (28%) | 4 (18%) | 11 (35%) |  |
| 5-FU monotherapy | 7 (9.1%) | 7 (13%) | 5 (23%) | 2 (6.5%) |  |
| No postoperative chemotherapy | 24 (31%) | 17 (32%) | 9 (41%) | 8 (26%) |  |
| Unknown | 1 (1.3%) | 1 (1.9%) | 0 (0%) | 1 (3.2%) |  |
| <sup>1</sup>Median (Q1-Q3); n (%) |  |  |  |  |  |
| <sup>2</sup>Wilcoxon rank sum exact test; Fisher's exact test |  |  |  |  |  |

Data are presented as median (IQR) or n (%). The primary molecular cohort comprised patients with an available pretreatment FOLFIRINOX sensitivity classification and biopsy tumour cellularity $\geq 10\%$ (n=53). FFX-Sens (n=31) and FFX-Res (n=22) are mutually exclusive subgroups of the primary molecular cohort. Percentages are calculated within each column; where data were missing, the number of evaluable patients is indicated.
IQR, interquartile range; CA19-9, carbohydrate antigen 19-9; CAP, College of American Pathologists tumour regression grade; ypN, pathological nodal stage after neoadjuvant therapy; R0, microscopically margin-negative resection; R1, microscopically margin-positive resection; FFX, FOLFIRINOX; FFX-Sens, classified as FOLFIRINOX-sensitive; FFX-Res, not classified as FOLFIRINOX-sensitive; 5-FU, 5-fluorouracil. FOLFIRINOX-related postoperative chemotherapy included FOLFIRINOX or FOLFIRI; gemcitabine-related postoperative chemotherapy included gemcitabine with or without capecitabine.

Given the cohort size and number of events, the baseline multivariable models were restricted to three covariates: pretreatment FFX classification, baseline CA19-9 and baseline tumour size. Complete-case analyses included 50 patients, with 34 deaths for OS and 37 events for DFS. Baseline covariates were selected because the analysis evaluated information available before treatment. A separate complete-case multivariable model (n=50) included pretreatment FFX classification, resection margin (R1 versus R0), pathological nodal status (positive versus negative) and CAP tumour regression grade (per grade increase). Analyses used available data without imputation.

Paired binary classifications were compared using the exact McNemar test, and paired continuous or ordinal measurements were compared using the Wilcoxon signed-rank test. All tests were two-sided. No adjustment for multiple comparisons was applied to exploratory or supplementary analyses. Statistical analyses were performed using R version 4.6.1. No formal sample size calculation was performed; cohort size was determined by the number of patients with usable tissue in the source surgical cohort over the study period, rather than by a pre-specified power calculation.

## Results

### Pretreatment FOLFIRINOX sensitivity is associated with outcome after neoadjuvant FOLFIRINOX

Of 141 patients in the source surgical cohort, 113 met the clinical eligibility criteria; tissue availability yielded an overall translational cohort of 77 patients (Figure 1). Clinical and pathological characteristics are summarised in Table 1. Median follow-up was 89.1 months (95% CI 71.6-102.8).

A pretreatment transcriptomic FOLFIRINOX sensitivity classification was available for 60 patients. Using the previously established 10% tumour-cellularity threshold for molecular evaluability [14,15], the primary molecular cohort comprised 53 patients, of whom 31 were classified as FFX-Sens and 22 as FFX-Res (Figure 1).

Pretreatment FFX-Sens status was associated with longer OS than FFX-Res status (HR 0.48, 95% CI 0.25-0.93; p=0.029; Figure 2A). The association with DFS was not statistically significant (HR 0.60, 95% CI 0.32-1.12; p=0.107; Figure 2B). Median OS was 59.6 months in the FFX-Sens group and 20.5 months in the FFX-Res group; median DFS was 30.1 and 11.3 months, respectively. We next examined clinically relevant baseline factors. In univariable analyses, FFX status was the only factor significantly associated with OS; performance status showed a non-significant association with OS (HR 1.84, 95% CI 0.96-3.54; p=0.067) and was associated with DFS (HR 2.05, 95% CI 1.09-3.88; p=0.026; Supplementary Table S1). The complete-case multivariable models included FFX classification, baseline CA19-9 and baseline tumour size in 50 patients. FFX-Sens remained associated with longer OS (adjusted HR 0.47, 95% CI 0.23-0.96; p=0.037; Figure 2C), whereas the adjusted association with DFS was not statistically significant (adjusted HR 0.55, 95% CI 0.28-1.10; p=0.090; Figure 2D). The magnitude and direction of the FFX estimates changed little after adjustment, although the DFS estimate remained imprecise.

**Figure 2.**
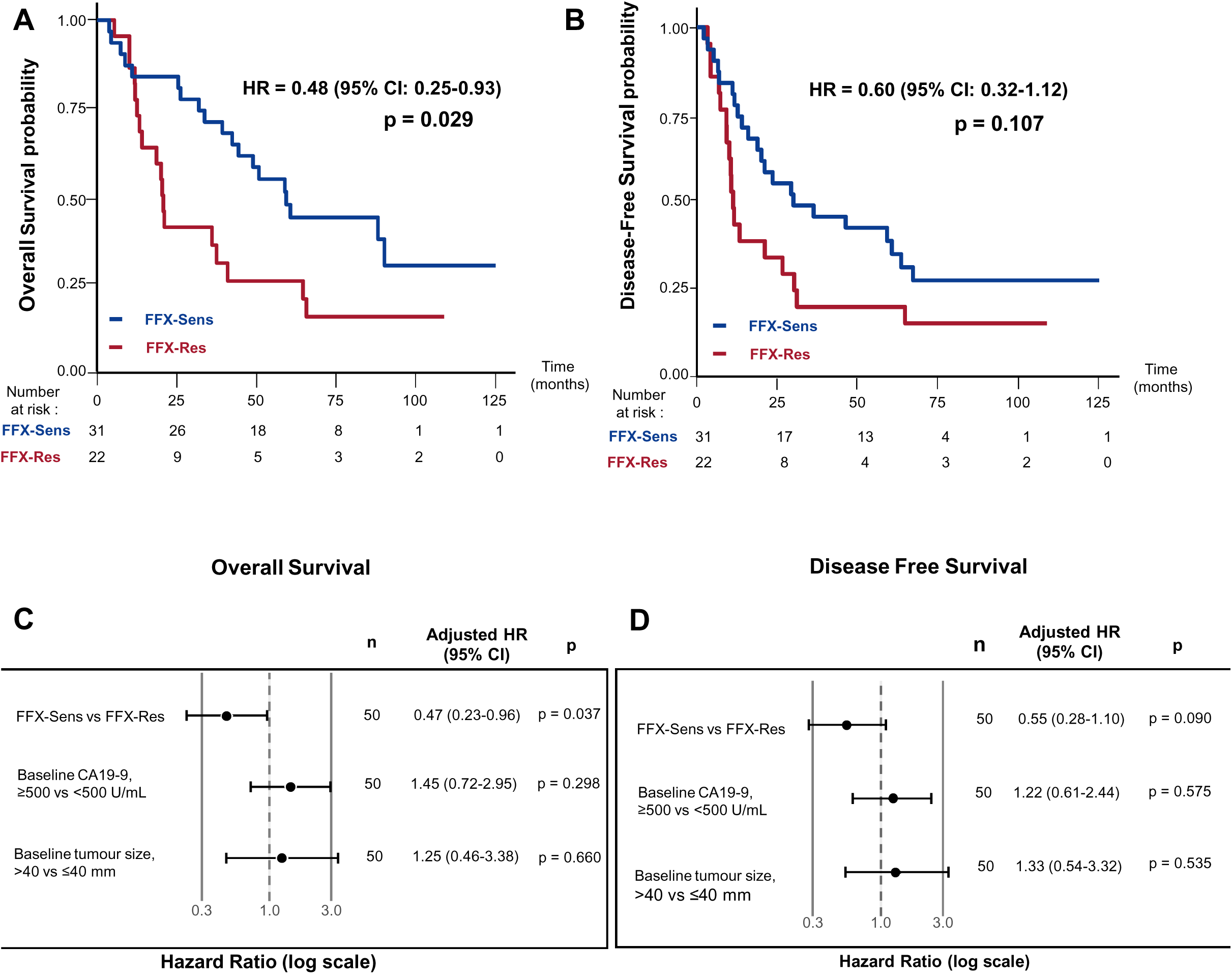
Association between pretreatment transcriptomic FOLFIRINOX classification and survival after neoadjuvant FOLFIRINOX. (A) Overall survival according to pretreatment FOLFIRINOX sensitivity. (B) Disease-free survival according to pretreatment FOLFIRINOX sensitivity. Hazard ratios and p-values in panels A and B were derived from univariable Cox proportional-hazards models. (C) Multivariable Cox regression analysis of overall survival and (D) Multivariable Cox regression analysis of disease-free survival. Both models included pretreatment FFX classification, baseline CA19-9 (≥500 versus <500 U/mL) and baseline tumour size (>40 versus ≤40 mm) and were restricted to 50 patients with complete data. Adjusted hazard ratios are shown with 95% confidence intervals. FFX, FOLFIRINOX; FFX-Sens, classified as FOLFIRINOX-sensitive; FFX-Res, not classified as FOLFIRINOX-sensitive; OS, overall survival; DFS, disease-free survival; HR, hazard ratio; CI, confidence interval; CA19-9, carbohydrate antigen 19-9.

### Residual tumours show directional reclassification of FOLFIRINOX sensitivity after treatment

The availability of paired pretreatment biopsies and surgical specimens provided an opportunity to determine whether the transcriptomic FOLFIRINOX sensitivity state remained stable during treatment. Longitudinal analysis was restricted to 35 patients with molecularly evaluable pretreatment and residual-tumour specimens.

A marked directional change in FOLFIRINOX classification was observed after treatment. Among 24 tumours classified as FFX-Sens before FOLFIRINOX, 15 were classified as FFX-Res in the residual tumour, whereas only 4 of the 11 pretreatment FFX-Res tumours changed to FFX-Sens. Nine tumours remained FFX-Sens and seven remained FFX-Res. The asymmetry between Sens-to-Res and Res-to-Sens transitions was statistically significant (exact McNemar p=0.019; Figure 3A).

**Figure 3.**
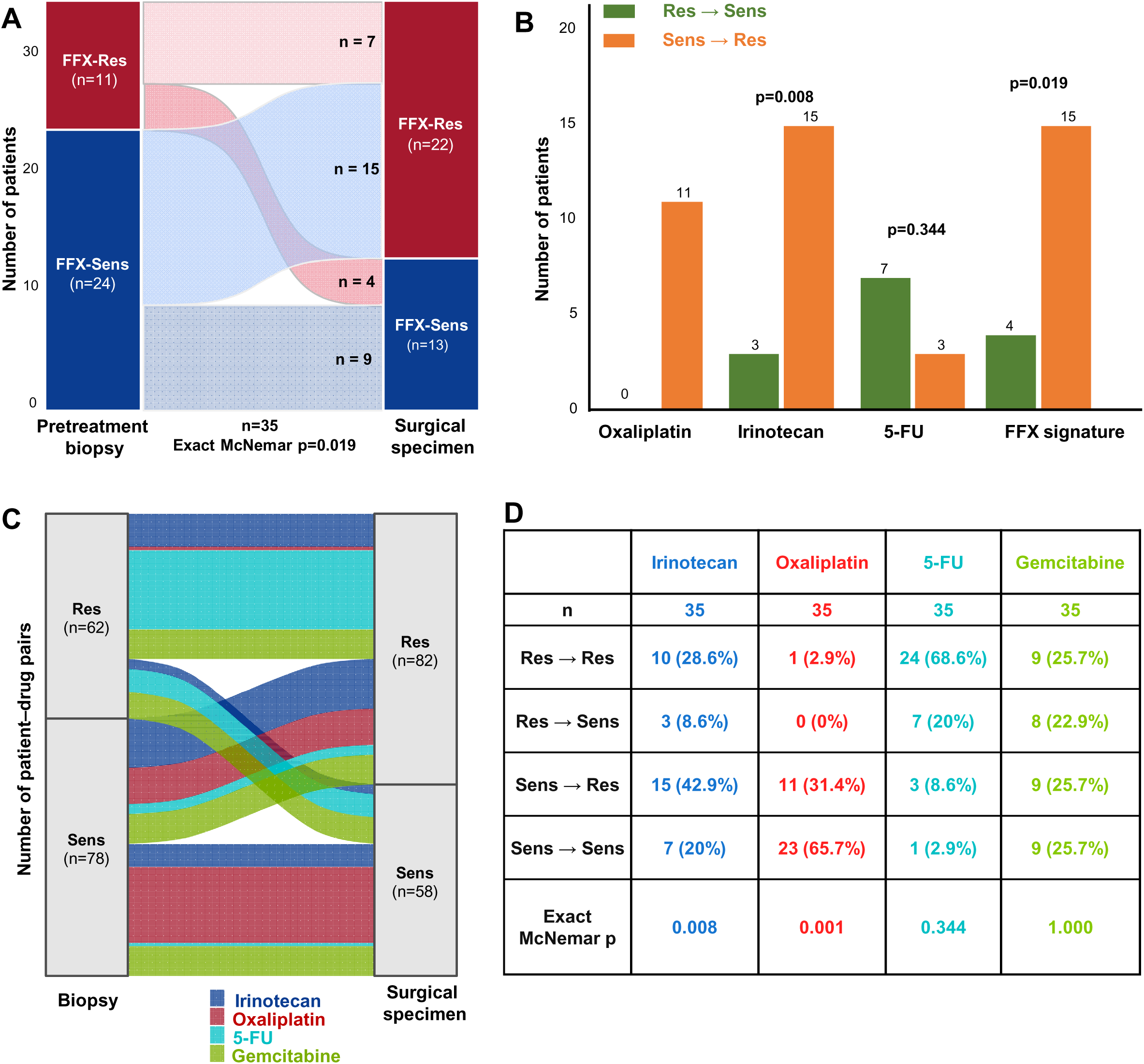
Directional reclassification of transcriptomic FOLFIRINOX sensitivity after neoadjuvant FOLFIRINOX. (A) Paired changes in FOLFIRINOX sensitivity classification between pretreatment biopsies and residual surgical tumours in the paired molecular cohort (n=35). (B) Direction of paired classification changes for the individual FOLFIRINOX components and the composite FFX classification. (C) Directional reclassification of sensitivity to each individual drug between pretreatment biopsy and surgical specimen (35 patients; 140 paired patient-drug classifications), including gemcitabine, which is not a component of the FOLFIRINOX regimen and was included as an exploratory negative control. (D) Exact transition counts (Res→Res, Res→Sens, Sens→Res, Sens→Sens) and McNemar p-values corresponding to panel C, for each of the four drugs. FFX, FOLFIRINOX; FFX-Sens, classified as FOLFIRINOX-sensitive; FFX-Res, not classified as FOLFIRINOX-sensitive; Sens, classified as sensitive; Res, not classified as sensitive; 5-FU, 5-fluorouracil.

Drug-specific analyses showed directional shifts towards classification as Res for irinotecan (15 Sens→Res versus 3 Res→Sens; exact McNemar p=0.008) and oxaliplatin (11 Sens→Res versus 0 Res→Sens; exact McNemar p=0.001), whereas 5-fluorouracil showed no comparable directional change (3 Sens→Res versus 7 Res→Sens; exact McNemar p=0.344; Figure 3B). GemCore was examined as an exploratory negative control because gemcitabine is not a component of FOLFIRINOX and showed no directional change (9 Sens→Res versus 8 Res→Sens; exact McNemar p=1.000; Figure 3C,D). Full transition counts and exact McNemar p-values for all four drugs are provided in Figure 3D.

To determine whether this directional change in FOLFIRINOX sensitivity occurred in the context of a broader shift in PDAC molecular phenotype, we examined the basal/classical status using PAMG and PurIST as exploratory secondary classifiers [10,23]. Among all patients with paired PAMG measurements (n=56), PAMG scores decreased after treatment from a median of 107 to 90 (paired Wilcoxon p=0.035). When restricted to the paired molecular cohort (n=35), the same direction was maintained, although the difference was attenuated and no longer statistically significant (median 99 versus 78; p=0.088). PurIST classification was comparatively stable in these 35 paired tumours: 30 tumours were classified as classical and five as basal-like before treatment, compared with 28 classical and seven basal-like residual tumours (exact McNemar p=0.727; Supplementary Figure S1). Thus, directional FOLFIRINOX reclassification was observed alongside a modest exploratory shift in PAMG, but without a significant change in PurIST subtype classification.

These paired analyses therefore indicate that the transcriptomic FOLFIRINOX sensitivity classification is not necessarily preserved in residual tumours after treatment. The observed changes are compatible with treatment-associated directional reclassification, but the study design cannot distinguish selection of pre-existing tumour populations from transcriptional adaptation, spatial heterogeneity or other sources of longitudinal variation. Accordingly, these findings should not be interpreted as direct evidence of acquired resistance.

### Pretreatment molecular sensitivity remains associated with outcome after adjustment for standard postoperative pathological factors

Major pathological responses were uncommon, with CAP tumour regression grades 0-1 observed in 6 of 72 evaluable surgical specimens (8.3%). Major responses occurred in both FFX-Sens and FFX-Res tumours, and pretreatment FFX status was not significantly associated with CAP tumour regression grade (Table 1), indicating that the pretreatment transcriptomic classification did not merely recapitulate the pathological response observed after FOLFIRINOX.

Major pathological response (CAP tumour regression grades 0-1) was associated with longer survival. No deaths occurred in the CAP 0-1 group during follow-up, precluding estimation of a Cox HR for OS; the Kaplan-Meier curves differed significantly (log-rank p=0.005; Figure 4A). CAP tumour regression grades 2-3 were associated with shorter DFS than CAP tumour regression grades 0-1 (HR 4.13, 95% CI 1.004-16.98; p=0.049; Figure 4B).

**Figure 4.**
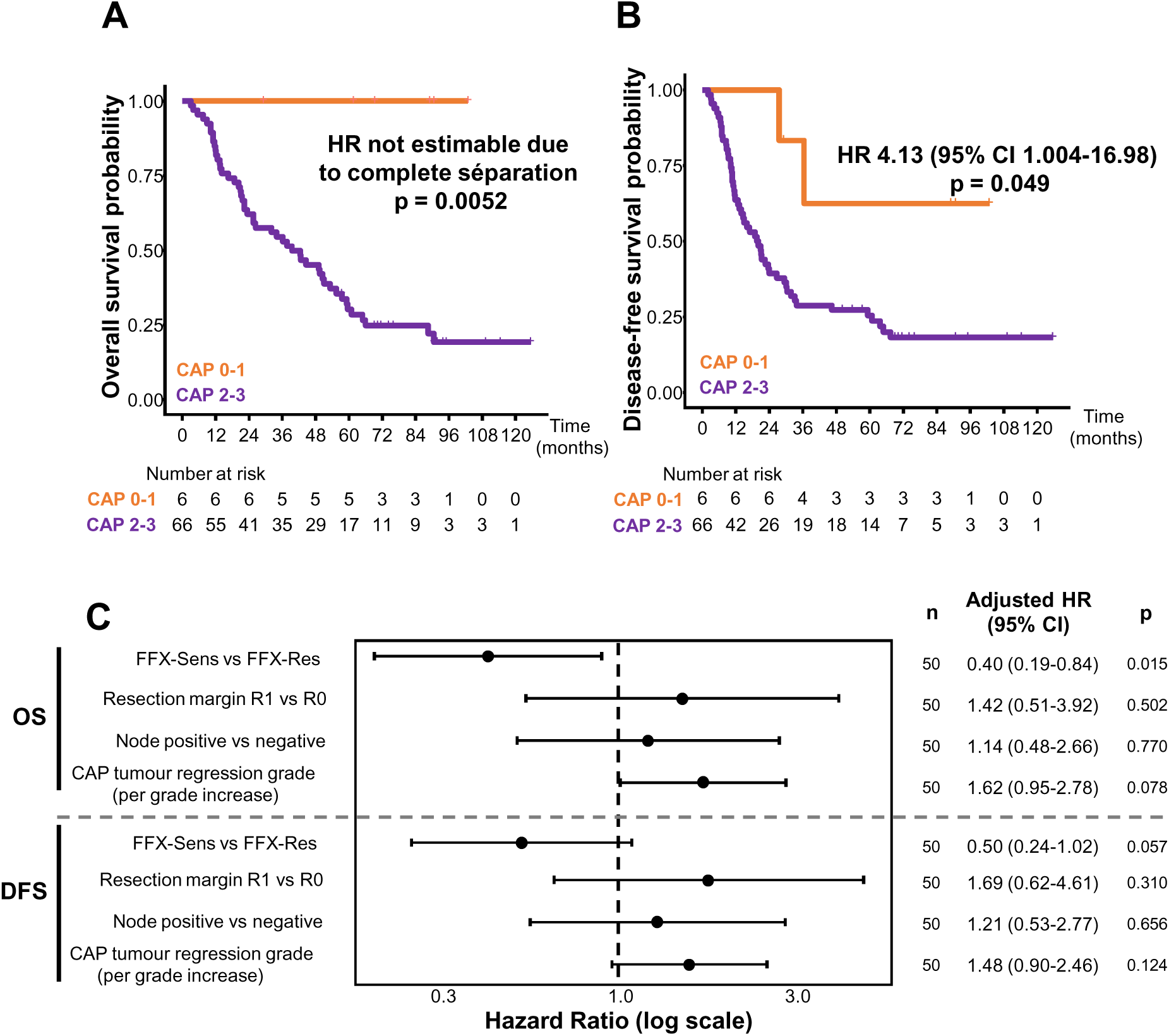
Association of CAP tumour regression grade and pretreatment FOLFIRINOX classification with survival. **(A)** Overall survival according to CAP tumour regression grade. **(B)** Disease-free survival according to CAP tumour regression grade. **(C)** Multivariable Cox analyses of overall survival and disease-free survival included pretreatment FFX classification, resection margin, pathological nodal status and CAP tumour regression grade and were restricted to 50 patients with complete data. FFX, FOLFIRINOX; FFX-Sens, classified as FOLFIRINOX-sensitive; FFX-Res, not classified as FOLFIRINOX-sensitive; OS, overall survival; DFS, disease-free survival; HR, hazard ratio; CI, confidence interval; CAP, College of American Pathologists.

To determine whether the pretreatment classifier remained informative after accounting for standard postoperative pathological factors, these factors were first examined individually. R1 resection margin was associated with shorter OS and DFS in univariable analyses, whereas pathological nodal status and CAP tumour regression grade did not reach statistical significance for either endpoint (Supplementary Table S2). A multivariable Cox model then included FFX classification, resection margin, pathological nodal status and CAP tumour regression grade (n=50). FFX-Sens status remained associated with longer OS (adjusted HR 0.40, 95% CI 0.19-0.84; p=0.015) but was not significantly associated with DFS (adjusted HR 0.50, 95% CI 0.24-1.02; p=0.057; Figure 4C). In the adjusted models, none of the postoperative pathological factors reached statistical significance. Full univariable and multivariable estimates are provided in Supplementary Table S2.

## Discussion

In this study, previously developed transcriptomic signatures of sensitivity to the individual components of FOLFIRINOX were evaluated in patients with PDAC treated with neoadjuvant FOLFIRINOX followed by surgical resection. Three findings emerge. First, pretreatment FFX-Sens status was associated with longer OS, whereas the DFS association was directionally concordant but not statistically significant. Second, the association with OS persisted after adjustment for baseline CA19-9 and tumour size and, in a separate model, for postoperative pathological factors. Third, paired analysis showed a significant directional shift from FFX-Sens towards FFX-Res after treatment. Together, these findings extend a clinically evaluated drug-sensitivity framework to the neoadjuvant BR/LA setting and, importantly, provide the first longitudinal evidence that FOLFIRINOX sensitivity classification may not be preserved in residual disease after treatment.

The pretreatment findings extend previous evaluations of this transcriptomic strategy in resected and advanced PDAC [12–18] to patients receiving FOLFIRINOX before surgery. To our knowledge, this is the first application of these classifiers in the neoadjuvant BR/LA setting. This distinction is clinically important because treatment selection is made before tumour sensitivity is known, and failure of induction therapy may preclude curative-intent resection. In the recent TAPS cohort of 1,835 patients receiving initial (m)FOLFIRINOX, resection was ultimately performed in 53.1% of patients with borderline-resectable and 17.8% of those with locally advanced disease, with baseline stage, CA19-9, performance status and tumour size associated with the probability of resection [24]. These clinical variables estimate resectability but do not directly measure tumour-specific drug sensitivity. In our cohort, the association between pretreatment FFX sensitivity and OS changed little after adjustment for baseline CA19-9 and tumour size. However, because all patients received the same neoadjuvant regimen, the study cannot distinguish a treatment-specific predictive effect from a prognostic association. Postoperative chemotherapy was also not standardised and may have contributed to long-term outcome. The findings should therefore be interpreted as outcome stratification under FOLFIRINOX exposure rather than proof of a specific benefit from neoadjuvant FOLFIRINOX.

Pathological response after neoadjuvant therapy is an established prognostic factor in PDAC [27,28]. In the separate postoperative model, pretreatment FFX sensitivity remained associated with OS after adjustment for resection margin, nodal status and CAP tumour regression grade. Because nodal status and CAP tumour regression grade were not significantly associated with outcome in this small selected cohort, this analysis should be regarded as supportive rather than as definitive evidence of incremental prognostic value.

A key strength of the study is the paired design, which examines the molecular state of the tumour that persists after FOLFIRINOX. The predominance of FFX-Sens-to-FFX-Res transitions indicates directional reclassification during treatment, with the strongest shifts involving irinotecan and oxaliplatin, whereas 5-fluorouracil showed no comparable directional change. GemCore, examined as an exploratory negative control because gemcitabine was not administered, also showed no directional shift. The modest decrease in PAMG score and relative stability of PurIST classification further suggest that FOLFIRINOX reclassification cannot simply be reduced to a uniform classical-to-basal subtype conversion. Previous studies have demonstrated substantial transcriptional remodelling of PDAC after neoadjuvant treatment [29–31]. To our knowledge, however, the present study is the first matched analysis applying validated drug-specific FOLFIRINOX sensitivity classifiers to pretreatment and residual-tumour specimens from the same patients. The finding that drug-sensitivity classification is not necessarily preserved raises the hypothesis that reassessment of residual disease could provide information not available from the diagnostic biopsy alone. Whether such reassessment has prognostic or treatment-selection value cannot be determined from the present study. Moreover, the observed transitions may reflect selection of pre-existing tumour populations, transcriptional adaptation, spatial heterogeneity or a combination of these mechanisms and should therefore not be interpreted as direct evidence of acquired resistance.

Whether the reclassification of FOLFIRINOX sensitivity observed in residual disease could inform postoperative treatment remains an open question. PANACHE 02 (PRODIGE93, NCT07044453) is already evaluating risk-adapted adjuvant chemotherapy based on post-neoadjuvant pathological staging [32], illustrating the clinical relevance of reassessing tumour status after neoadjuvant therapy. Whether molecular reassessment could provide complementary information for postoperative treatment selection warrants prospective investigation.

The ongoing PRODIGE 104A-NEOPREDICT trial is prospectively testing transcriptomic-guided neoadjuvant treatment selection in borderline-resectable PDAC [33]. This strategy is already being prospectively evaluated in metastatic PDAC through PACsign-01 and GemSign-01 [25,26], whereas NEOPREDICT extends transcriptomic-guided chemotherapy selection to the potentially curative neoadjuvant setting. Patients with a GEM-positive transcriptomic signature are randomised to gemcitabine/nab-paclitaxel or mFOLFIRINOX, thereby testing the principle of using a pretreatment transcriptomic signature to inform regimen selection at the neoadjuvant decision point. NEOPREDICT does not directly validate the FOLFIRINOX-component classifiers evaluated here. The present retrospective data are therefore complementary: they support prospective evaluation of FOLFIRINOX sensitivity classification in the neoadjuvant setting and show that the drug-sensitivity state measured before treatment may not be preserved in residual disease.

Several limitations should be considered. The study was retrospective, single-centre and included a relatively small number of patients. Multivariable analyses were deliberately parsimonious. The baseline models included only FFX classification, CA19-9 and tumour size and were restricted to 50 complete cases, with 34 deaths and 37 DFS events. These models should therefore be considered exploratory and not as comprehensive clinical prognostic models. Importantly, only patients who ultimately underwent surgery were included; patients who progressed, died or became unresectable during neoadjuvant treatment were not represented. This creates substantial selection bias towards patients who successfully completed the neoadjuvant-to-surgery pathway and prevents evaluation of the classifier in the full population initiating neoadjuvant FOLFIRINOX. This selection is particularly relevant to the clinical interpretation of the classifier because the present study cannot determine whether pretreatment FFX classification predicts failure to reach surgery, one of the key outcomes for a biomarker intended to guide neoadjuvant treatment selection. The additional exclusion of patients with recurrence or death within 3 months after surgery may also have favoured longer-surviving patients and limits interpretation of absolute survival estimates. Generalisability is therefore limited to patients who reach resection after neoadjuvant FOLFIRINOX in a single-centre setting. The absence of an alternative neoadjuvant treatment group also prevents formal demonstration of treatment-specific predictive value. The paired analysis was restricted to 35 molecularly evaluable cases and cannot distinguish biological evolution from spatial sampling heterogeneity.

In conclusion, pretreatment transcriptomic FOLFIRINOX sensitivity was associated with OS in this selected cohort of resected BR/LA PDAC treated with neoadjuvant FOLFIRINOX, including after adjustment for selected baseline and postoperative factors. The association with DFS was not statistically significant. Paired analysis showed directional reclassification of FOLFIRINOX sensitivity in residual disease. These findings extend the evaluation of this transcriptomic strategy to the neoadjuvant setting and provide a retrospective rationale for prospective studies testing whether pretreatment classification can improve regimen selection and whether reassessment of residual disease can inform subsequent therapeutic decisions.

## Data availability

The clinical and transcriptomic data generated and/or analysed during the current study are available from the corresponding author on reasonable request, subject to applicable ethical, patient-confidentiality and data-protection restrictions. The FOLFIRINOX transcriptomic sensitivity classifiers used in this study are covered by intellectual property rights and are not publicly available. Access to proprietary classifier components may be subject to additional licensing restrictions.

## Author contributions

N.D. conceived the study, defined the translational and analytical strategy, supervised the project, contributed to data analysis and interpretation, and co-wrote the original manuscript with C.G. C.G. curated the clinical data, performed the analyses, interpreted the results, prepared the figures, and co-wrote the original manuscript. N.F. developed and implemented the transcriptomic and computational analyses and contributed to data interpretation and visualisation. B.C. contributed to the study design, clinical and statistical interpretation of the results, and co-supervised the study. M.R. and J.R. provided technical assistance for sample processing and histopathological analyses. F.P. and M.K. contributed pathological expertise, sample assessment and pathological annotation. J.G. contributed surgical samples and associated clinical data. P.R. contributed to the analysis and interpretation of the results and reviewed and edited the manuscript. A.B. contributed to sample collection and the analysis and interpretation of the results. All authors critically reviewed the manuscript, approved the final version, and agree to be accountable for the work.

## Declaration of Generative AI and AI-assisted technologies in the writing process

During the preparation of this work, the authors used ChatGPT (OpenAI) to assist with language editing, structural consistency and compliance checks. After using this tool, the authors reviewed and edited the content as needed and take full responsibility for the content of the publication.

**Supplementary Figure S1.**

**Longitudinal changes in PDAC molecular phenotype after neoadjuvant FOLFIRINOX.**

**(A)** PAMG scores in paired pretreatment biopsies and residual surgical tumours.
**(B)** PurIST classification in the paired molecular cohort (n=35).

PDAC, pancreatic ductal adenocarcinoma; PAMG, pancreatic adenocarcinoma molecular gradient; PurIST, Purity Independent Subtyping of Tumors.

## Abbreviations

BR/LA: borderline resectable/locally advanced
FFX: FOLFIRINOX
FFX-Sens: classified as FOLFIRINOX-sensitive
FFX-Res: not classified as FOLFIRINOX-sensitive
PAMG: pancreatic adenocarcinoma molecular gradient
PurIST: Purity Independent Subtyping of Tumors

