## Supplementary Figure 1 for "Transcriptomic FOLFIRINOX sensitivity signatures stratify overall survival and reveal directional reclassification after neoadjuvant FOLFIRINOX in borderline resectable and locally advanced pancreatic cancer"

### Slide 1
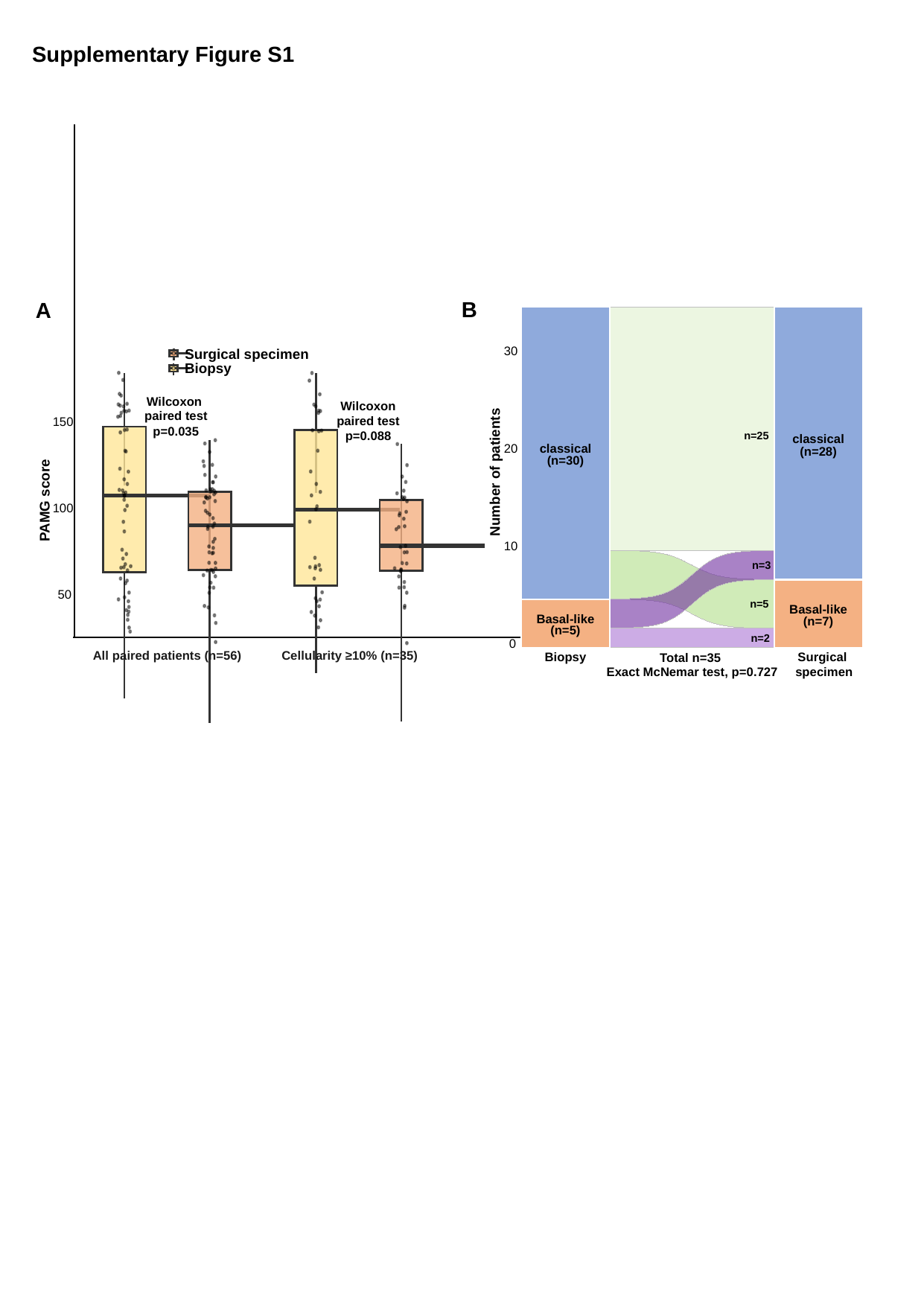

Supplementary Figure S1
B
A
30
classical
classical
20
(n=28)
(n=30)
Number of patients
10
Basal-like
Basal-like
(n=7)
(n=5)
0
Surgical
specimen
Total n=35
Exact McNemar test, p=0.727
Biopsy
n=25
n=3
n=5
n=2
Surgical specimen
Biopsy
Wilcoxon
paired test
p=0.088
Wilcoxon
paired test
p=0.035
150
PAMG score
100
50
Cellularity ≥10% (n=35)
All paired patients (n=56)
