## Supplementary Table 1 for "Transcriptomic FOLFIRINOX sensitivity signatures stratify overall survival and reveal directional reclassification after neoadjuvant FOLFIRINOX in borderline resectable and locally advanced pancreatic cancer"

### Supplementary Table S1. Univariable and multivariable Cox proportional-hazards analyses for overall and disease-free survival in the primary molecular cohort

### A. Overall survival

| **Variable** | **n (univariable)** | **Events**  **(univariable)** | **Univariable HR (95% CI)** | **p** | **Adjusted HR (95% CI)** | **p** |
| --- | --- | --- | --- | --- | --- | --- |
| **FFX-Sens versus FFX-Res** | 53 | 37 | **0.48 (0.25-0.93)** | **0.029** | **0.47 (0.23-0.96)** | **0.037** |
| **Baseline CA19-9, ≥ 500 versus < 500 U/mL** | 52 | 36 | 1.48 (0.74-2.96) | 0.272 | 1.45 (0.72-2.95) | 0.298 |
| **Baseline tumour size, > 40 versus ≤ 40 mm** | 51 | 35 | 0.95 (0.37-2.45) | 0.912 | 1.25 (0.46-3.38) | 0.660 |
| **Age, > 65 versus ≤ 65 years** | 53 | 37 | 0.98 (0.51-1.87) | 0.949 |  |  |
| **Male versus Female** | 53 | 37 | 1.22 (0.63-2.36) | 0.551 |  |  |
| **Performance status, 1 versus 0** | 53 | 37 | 1.84 (0.96-3.54) | 0.067 |  |  |
| **Locally advanced versus borderline resectable** | 53 | 37 | 0.79 (0.34-1.80) | 0.570 |  |  |
| **Other tumour location versus head** | 53 | 37 | 1.46 (0.76-2.79) | 0.252 |  |  |

### B. Disease-free survival

| **Variable** | **n (univariable)** | **Events**  **(univariable)** | **Univariable HR (95% CI)** | **p** | **Adjusted HR (95% CI)** | **p** |
| --- | --- | --- | --- | --- | --- | --- |
| **FFX-Sens versus FFX-Res** | 53 | 40 | 0.60 (0.32-1.12) | 0.107 | 0.55 (0.28-1.10) | 0.090 |
| **Baseline CA19-9, ≥ 500 versus <500 U/mL** | 52 | 39 | 1.26 (0.64-2.49) | 0.509 | 1.22 (0.61-2.44) | 0.575 |
| **Baseline tumour size, > 40 versus ≤ 40 mm** | 51 | 38 | 1.11 (0.46-2.67) | 0.812 | 1.33 (0.54-3.32) | 0.535 |
| **Age, > 65 versus ≤ 65 years** | 53 | 40 | 0.94 (0.50-1.74) | 0.837 |  |  |
| **Male versus Female** | 53 | 40 | 1.23 (0.65-2.32) | 0.517 |  |  |
| **Performance status, 1 versus 0** | 53 | 40 | 2.05 (1.09-3.88) | 0.026 |  |  |
| **Locally advanced versus Borderline resectable** | 53 | 40 | 0.65 (0.29-1.47) | 0.302 |  |  |
| **Other tumour location versus head** | 53 | 40 | 1.39 (0.74-2.61) | 0.304 |  |  |

HR, hazard ratio; CI, confidence interval; CA19-9, carbohydrate antigen 19-9; FFX-Sens, classified as FOLFIRINOX-sensitive; FFX-Res, not classified as FOLFIRINOX-sensitive. Univariable analyses used available data without imputation. Both adjusted models included FFX classification, baseline CA19-9 and baseline tumour size and were restricted to 50 patients with complete data.
