## Supplementary table 2 for "Transcriptomic FOLFIRINOX sensitivity signatures stratify overall survival and reveal directional reclassification after neoadjuvant FOLFIRINOX in borderline resectable and locally advanced pancreatic cancer"

**Supplementary Table S2. Univariable and multivariable Cox proportional hazards analyses of pretreatment FOLFIRINOX classification and post-surgical pathological factors for overall and disease-free survival.**

A. **Overall Survival**

| **Variable** | **n**  **(Univariable)** | **Univariable HR (95% CI)** | **p** | **Adjusted HR (95% CI)** | **p** |
| --- | --- | --- | --- | --- | --- |
| FFX-Sens versus FFX-Res | 53 | **0.48 (0.25, 0.93)** | **0.029** | **0.40 (0.19, 0.84)** | **0.015** |
| Resection margin R1 versus R0 | 53 | **2.67 (1.10, 6.45)** | **0.029** | 1.42 (0.51, 3.92) | 0.502 |
| Node positive versus negative | 53 | 1.92 (0.95, 3.89) | 0.071 | 1.14 (0.48, 2.66) | 0.770 |
| CAP tumour regression grade (per grade increase) | 50 | 1.54 (0.94, 2.52) | 0.089 | 1.62 (0.95, 2.78) | 0.078 |

B. **Disease-free survival**

| **Variable** | **n (univariable)** | **Univariable HR (95% CI)** | **p** | **Adjusted HR (95% CI)** | **p** |
| --- | --- | --- | --- | --- | --- |
| FFX-Sens versus FFX-Res | 53 | 0.60 (0.32- 1.12) | 0.107 | 0.50 (0.24-1.02) | 0.057 |
| Resection margin R1 versus R0 | 53 | 2.87 (1.17- 7.00) | 0.021 | 1.69 (0.62-4.61) | 0.310 |
| Node positive versus negative | 53 | 1.85 (0.94- 3.65) | 0.075 | 1.21 (0.53-2.77) | 0.656 |
| CAP tumour regression grade (per grade increase) | 50 | 1.45 (0.93-2.26) | 0.102 | 1.48 (0.90-2.46) | 0.124 |

HR, hazard ratio; CI, confidence interval; CAP, College of American Pathologists; FFX-Sens, classified as FOLFIRINOX-sensitive; FFX-Res, not classified as FOLFIRINOX-sensitive; R0, microscopically margin-negative resection; R1, microscopically margin-positive resection. The n column indicates the number of patients included in each univariable analysis. Univariable analyses used available data without imputation. Both multivariable models included FFX classification, resection margin, pathological nodal status and CAP tumour regression grade and were restricted to 50 patients with complete data.
